# West Nile virus-inclusive single-cell RNA sequencing reveals heterogeneity in the type I interferon response within single cells

**DOI:** 10.1101/434852

**Authors:** Justin T. O’Neal, Amit A. Upadhyay, Amber Wolabaugh, Nirav B. Patel, Steven E. Bosinger, Mehul S. Suthar

**Author notes:** Address correspondence to Mehul S. Suthar,.

## Abstract

West Nile virus (WNV) is a neurotropic mosquito-borne flavivirus of global importance. Neuroinvasive WNV infection results in encephalitis and can lead to prolonged neurological impairment or death. Type I interferon (IFN-I) is crucial for promoting antiviral defenses through the induction of antiviral effectors, which function to restrict viral replication and spread. However, our understanding of the antiviral response to WNV infection is mostly derived from analysis of bulk cell populations. It is becoming increasingly apparent that substantial heterogeneity in cellular processes exists among individual cells, even within a seemingly homogenous cell population. Here, we present WNV-inclusive single-cell RNA sequencing (scRNA-seq), an approach to examine the transcriptional variation and viral RNA burden across single cells. We observed that only a few cells within the bulk population displayed robust transcription of IFN-β mRNA, and this did not appear to depend on viral RNA abundance within the same cell. Furthermore, we observed considerable transcriptional heterogeneity in the IFN-I response, with genes displaying high unimodal and bimodal expression patterns. Broadly, IFN-stimulated genes negatively correlated with viral RNA abundance, corresponding with a precipitous decline in expression in cells with high viral RNA levels. Altogether, we demonstrated the feasibility and utility of WNV-inclusive scRNA-seq as a high-throughput technique for single-cell transcriptomics and WNV RNA detection. This approach can be implemented in other models to provide insights into the cellular features of protective immunity and identify novel therapeutic targets.

**IMPORTANCE:** West Nile virus (WNV) is a clinically relevant pathogen responsible for recurrent epidemics of neuroinvasive disease. Type I interferon is essential for promoting an antiviral response against WNV infection; however, it is unclear how heterogeneity in the antiviral response at the single-cell level impacts viral control. Specifically, conventional approaches lack the ability to distinguish differences across cells with varying viral abundance. The significance of our research is to demonstrate a new technique for studying WNV infection at the single-cell level. We discovered extensive variation in antiviral gene expression and viral abundance across cells. This protocol can be applied to primary cells or *in vivo* models to better understand the underlying cellular heterogeneity following WNV infection for the development of targeted therapeutic strategies.

## INTRODUCTION

Mosquito-borne flaviviruses represent a significant public health burden, annually accounting for millions of infections worldwide that, in certain cases, can culminate in severe systemic or neuropathological outcomes (1–4). West Nile virus (WNV), a member of the *Flaviviridae* family, causes yearly epidemics of encephalitis and virus-induced myelitis on a global scale with nearly 50,000 reported cases of WNV disease and over 21,000 cases of neuroinvasive disease from 1999 to 2016 in the United States alone (1–4). Currently, there are no licensed vaccines or approved targeted therapeutics to prevent or treat WNV-infected patients, underscoring the need to better understand the cellular response to WNV infection (1–4).

Type I IFN (IFN-α/β or IFN-I) is the first line of defense against viral infection and coordinates the early antiviral programs to restrict viral replication, as well as shape the adaptive immune response (5–14). Loss of IFN-I signaling in WNV-infected mice results in uncontrolled viral replication and rapid mortality, demonstrating that the IFN-I response is required for protective immunity (9–11, 14, 15). Pattern recognition receptors, including toll-like receptors (TLRs) and retinoic acid-inducible gene I (RIG-I)-like receptors (RLRs), detect broad viral signatures, such as 5’-triphosphate ssRNA or dsRNA, in the cytosolic and endosomal compartments (9–11, 12, 14). For flavivirus infection, RLRs are critical for inducing IFN-I and binding to cytosolic viral RNA signals through adaptor proteins, such as mitochondrial antiviral signaling protein (MAVS), to activate transcription factors and induce interferon regulatory factor (IRF)-mediated transcription of IFN-β (*Ifnb1*) and a subset of IFN-stimulated genes (ISGs) (9, 11, 12, 14, 16–22). Signaling in both an autocrine and paracrine manner, secreted IFN-β binds IFN-I receptor (IFNAR1/2 heterodimer) to activate Janus kinases, Jak1 and Tyk2, which phosphorylate signal transducer and activator of transcription 1 (STAT1) and STAT2 (7, 18–21, 23–27). Phosphorylated STAT1 and STAT2 form a heterodimer and recruit IRF9 to form the ISG factor 3 (ISGF3) complex. The ISGF3 complex then translocates to the nucleus and induces IFN-stimulated response element (ISRE)-regulated genes, thereby reshaping the cellular landscape to an antiviral state (5–7, 9, 23–28).

The induction of IFN-I and ISGs within a bulk population of infected cells has been well characterized. However, mean values obtained via conventional bulk assays mask transcriptional differences between infected and bystander cells and obscure any heterogeneity present within the infected population. Recently, single-cell studies have examined the heterogeneity across virally infected cells. Findings with influenza virus, poliovirus, dengue virus (DENV) and Zika virus (ZIKV) have revealed extensive variation in viral RNA abundance within single cells (29–31). Using high dimensional mass cytometry by time-of-flight (CyTOF) analysis, others have described differences in IFN-induced and pro-inflammatory cytokine production in infected and bystander human dendritic cells following DENV infection (32). Studies examining IFN-I induction at the single-cell level have used fluorescently-tagged cells, single-mRNA molecule *in situ* hybridization, single-cell quantitative PCR (qPCR), and single-cell RNA sequencing (scRNA-seq) (16–19, 27, 33). Previous studies have found that only a small fraction of infected cells express *Ifnb1* mRNA (17–19, 27). This is thought to be attributable to stochasticity in signaling components and downstream signaling cascades leading to transcription factor activation or variability in the processes of *Ifnb1* expression, perhaps at the level of chromatin organization (16–19, 34–36). Using PRR agonists or nonproductive viral infection, others have demonstrated that IFN-I-dependent paracrine signaling is pivotal in amplifying the host antiviral response (16–19, 26, 27). Lastly, single-cell transcriptomic studies have also been used to globally investigate virus-host interactions and identify novel candidate genes for host-targeted therapeutics (31). Knockdown screens or knockout studies can only probe a subset of nonessential host genes, limiting their scope (37–42). However, virus-inclusive scRNA-seq is a powerful platform for the discovery of novel proviral and antiviral candidate genes in an unbiased manner as recently highlighted by Zanini and colleagues with DENV and ZIKV (31).

Altogether, these studies have shed considerable light on the transcriptional differences present in single cells, and specifically with *Ifnb1* expression and viral RNA abundance. However, we still lack a thorough understanding of the cellular heterogeneity in the IFN-I response following WNV infection. Population-level transcriptional analyses are valuable and widely used approaches, but in certain cases can belie gene expression patterns, such as bimodal variation, which can only be observed at single-cell resolution (18, 27, 33). To better understand the underlying transcriptional differences across cells with varying viral abundance, we developed WNV-inclusive scRNA-seq, a modified SMART-Seq protocol that incorporates a virus-specific primer for parallel recovery of host messenger RNA (mRNA) and viral RNA from single cells. We found that only a small fraction of cells exhibited robust *Ifnb1* expression, and this did not significantly correlate with high viral RNA. We observed considerable transcriptional heterogeneity in ISG expression and viral RNA abundance across cells. ISGs exhibited both unimodal and bimodal variation and were negatively correlated with intracellular viral RNA, displaying a steep decline in gene expression with increasing viral abundance. Combining single-cell mRNA sequencing with quantification of non-polyadenylated viral RNA, we present WNV-inclusive scRNA-seq as a high-throughput technique for single-cell transcriptome analysis of WNV-infected cells.

## RESULTS

### Population-level analysis of WNV infection in murine fibroblast L929 cells

We first modeled WNV infection kinetics in murine fibroblast L929 cells, an IFN-competent cell line extensively used to study IFN-I-dependent signaling (19, 43). Cells were infected at a multiplicity of infection (MOI) of 0.1, 1 or 10, as determined on BHK-21 cells, and intracellular viral envelope (E) protein immunostaining was performed at 6, 12, 24 and 48 hr post-infection. Infected cells were labeled with WNV E16 antibody (Ab), which recognizes a domain III (DIII) neutralizing epitope within the E protein (44). For all three MOIs, nearly 100% of cells stained positive for intracellular viral E protein by 48 hr post-infection (Fig. 1A). At a MOI of 10, intracellular viral E protein was detected in nearly 100% of cells as early as 24 hr post-infection, suggesting that the majority of these cells were likely infected during primary virus adsorption (Fig. 1A). For cells infected at a MOI of 1, intracellular viral E protein was detected in approximately 60% of cells at 24 hr post-infection (Fig. 1A).

**Figure 1.**
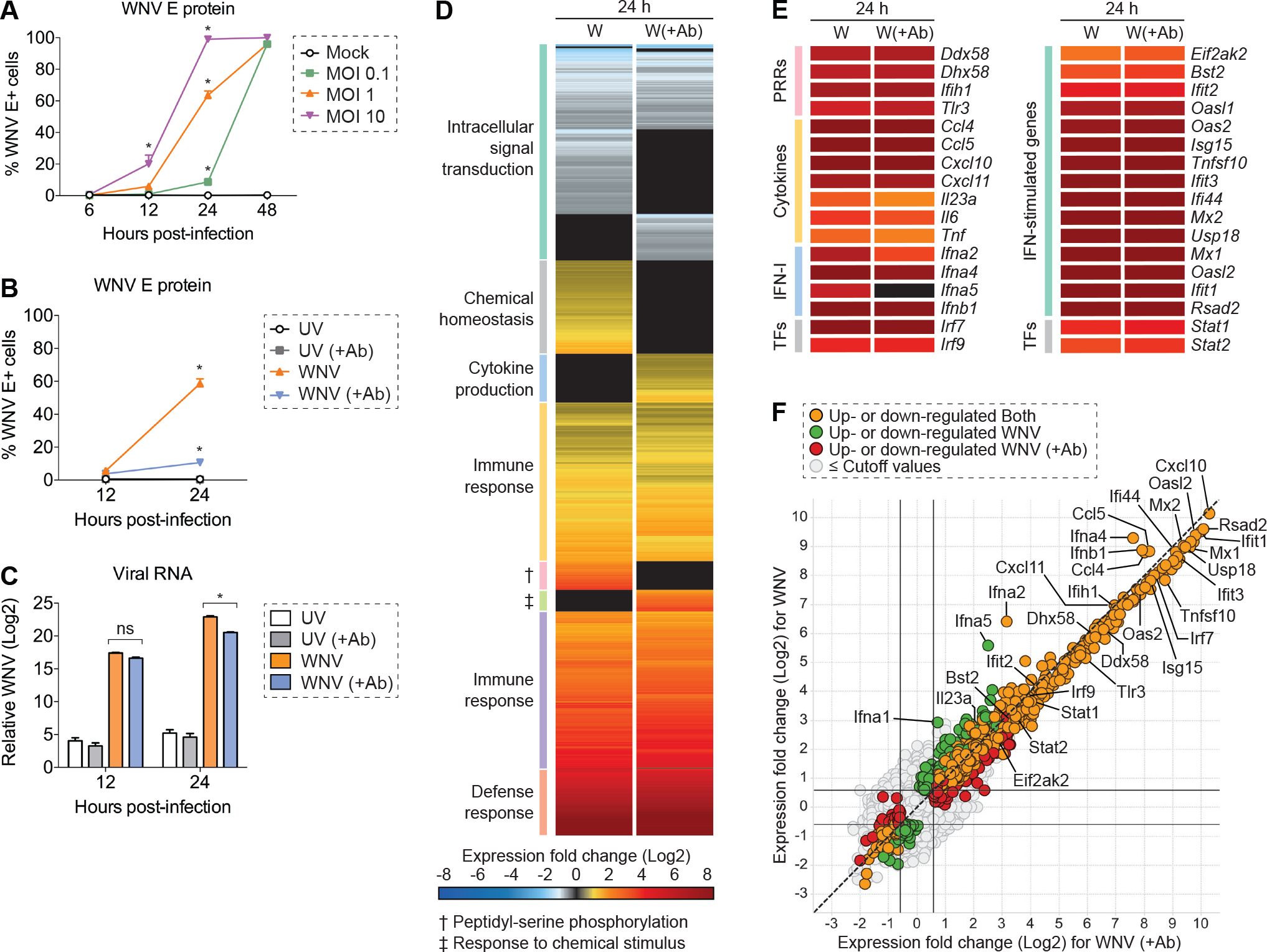
Population-level analysis of WNV infection in L929 cells. L929 cells were infected with WNV at a MOI of 0.1, 1, or 10 and incubated for 6, 12, 24, or 48 hr (n = 3). (A-B) Intracellular viral E protein staining was performed by flow cytometry using WNV E16 antibody. (B-C) L929 cells were inoculated with UV-inactivated WNV (UV) or WNV at a MOI of 1 and incubated for 12 or 24 hr in the presence (+Ab) or absence of WNV E16 neutralizing antibody (5 μg/mL) to reduce *in vitro* spread (n = 3). (C) Viral RNA quantification was measured by qPCR, and C_T_ values were normalized to the reference gene *Gapdh* and represented as fold change over time-matched mock values. (A-C) Two-way ANOVA with multiple comparison correction was used to test for significance (\**p* < 0.05). (D-F) Cells were infected as in and examined by bulk RNA-seq analysis at 24 hr post-infection (n = 3). (D-E) Heat map showing mean gene expression values normalized and represented as fold change over time-matched mock values. Expression fold change values correspond to the color gradient (bottom). (D) Gene cluster description can be found on the left. (E) Expression fold change displayed for a panel of select genes. (F) Scatter plot for comparison of up-regulated and down-regulated genes in WNV and WNV (+Ab) conditions. Cut-off values were as follows: 1.5 fold change and *p* < 0.01.

To diminish asynchronous second-round infection, cells were infected with WNV (MOI of 1) and incubated in the presence of WNV E16 neutralizing Ab. Inoculation with UV-inactivated WNV served as a non-replicating input control for internalized viral RNA, and no expression of viral E protein was detected (Fig. 1B, 1C). Notably, limiting *in vitro* spread resulted in a 5.5-fold decrease in the percentage of viral E protein-positive cells (10.6%) at 24 hr post-infection, corresponding with a comparable 5.2-fold reduction in viral RNA levels (Fig. 1B, 1C). Collectively, these two conditions, WNV and WNV (+Ab), provide a cell population with a range of viral abundance and another of predominantly bystander cells with which to survey the IFN-I response at the population and single-cell level in all subsequent analyses.

Before pursuing a single-cell approach, we next sought to evaluate transcriptional changes following WNV infection at the population level by bulk RNA-seq. As expected, numerous genes associated with the innate immune response and antiviral defense response were up-regulated following infection (Fig. 1D, 1E). Furthermore, the majority of these genes were expressed at similar levels independent of reduced asynchronous second-round infection (Fig. 1D, 1E). ISGs and PRR genes exhibited a more consistent level of mean gene expression across these two conditions (Fig. 1E). Conversely, IFN-I and cytokine genes displayed the most variability in expression between genes within their respective categories (Fig. 1E). Most notably, *Ifna2* and *Ifna5* displayed around two-fold higher levels of expression when allowing for *in vitro* spread, although *Ifna2* dropped outside of the pre-selected significance cutoff (*p* < 0.01; Fig. 1E, 1F). This population-level analysis provides a contextual fundamental framework from which to build as we examine the transcriptional differences observed across single cells. Leveraging single-cell sequencing techniques complemented with viral RNA detection, we next extended the resolution of our analysis to single cells to better understand the underlying transcriptional heterogeneity present following WNV infection.

### WNV-inclusive scRNA-seq captures mRNA and viral RNA from single cells

WNV-inclusive scRNA-seq is adapted from the well-established Smart-seq2 protocol (45) and the commercially available SMART-Seq v4 Ultra Low Input RNA Kit (Takara) used for scRNA-seq. The SMART-Seq v4 protocol is modified to include a virus-specific primer (WNV SC primer) during the reverse transcription (RT) step. For scRNA-seq analysis, L929 cells are inoculated with virus for 1 hr at a MOI of 1 and then incubated in the presence or absence of WNV E16 neutralizing Ab (44) for 24 hr (Fig. 2A). Viable single cells are sorted by conventional flow cytometry into 96-well plates containing 10 μL lysis buffer per well (Fig. 2B). In the RT reaction, 3’ SMART-Seq CDS Primer II A (30-nucleotide poly-dT sequence with a 5’ 25-nucleotide ISPCR universal anchor sequence (45)) and WNV SC primer (21-nucleotide sequence complementary to positive-strand viral RNA with a 5’ 25-nucleotide ISPCR universal anchor sequence (45)) are added to capture host transcripts and viral RNA, respectively (Fig. 2C). Following template switching, PCR Primer II A served as the primer for parallel downstream amplification of both host and viral complementary DNA (cDNA) (Fig. 2C). Samples underwent Nextera tagmentation and were sequenced on an Illumina HiSeq at a depth of approximately 1 million reads per cell (27, 46). Altogether, we successfully captured and profiled a total of 127 cells across three conditions: Mock, WNV, and WNV (+Ab). The outlined approach delivers exceptional coverage and sequencing depth allowing for accurate quantification of host transcripts and non-polyadenylated viral RNA.

**Figure 2.**
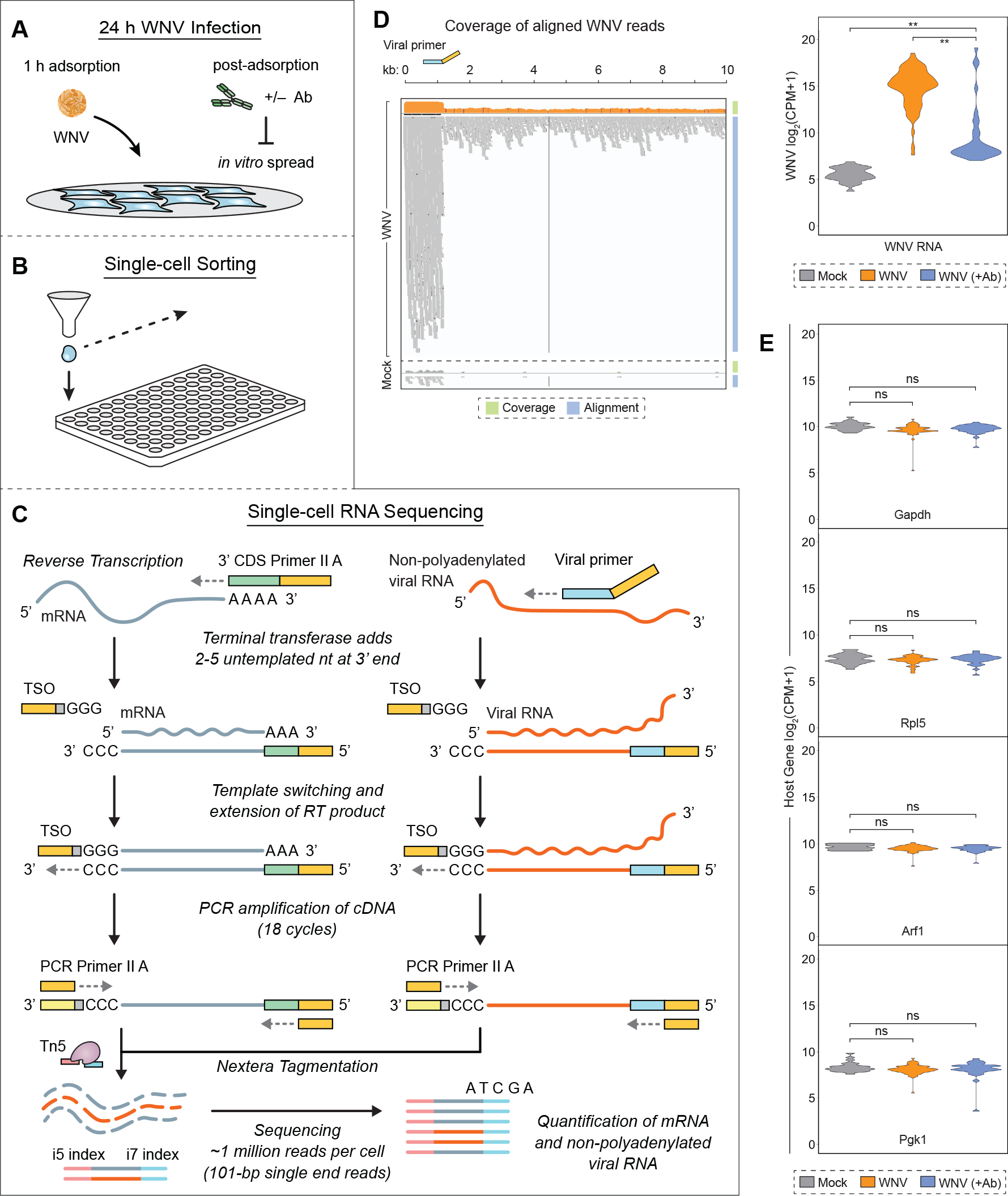
WNV-inclusive single-cell RNA sequencing. (A) L929 cells were infected with WNV (MOI of 1) and incubated in the presence of the WNV E16 neutralizing Ab (5 μg/mL) to limit *in vitro* spread. (B) Single cells were sorted into 96-well PCR plates containing 10 μL lysis buffer per well. (C) During reverse transcription (RT), 3’ SMART-Seq CDS Primer II A (30-nucleotide poly-dT sequence with a 5’ 25-nucleotide ISPCR universal anchor sequence) and a WNV-specific viral primer (21-nucleotide sequence complementary to positive-strand viral RNA with a 5’ 25-nucleotide ISPCR universal anchor sequence) are added to capture host transcripts and viral RNA, respectively. When the reverse transcriptase reaches the 5’ end of both host mRNA and viral RNA, its terminal transferase activity adds 2-5 untemplated nucleotides that serve as an anchor for the template-switching oligo (TSO), which allows extension of the RT product with sequence complementary to the universal anchor sequence. PCR Primer II A binds this sequence for concurrent amplification of both host and viral cDNA. In the final library preparation step, transposase 5 (Tn5)-based Nextera tagmentation adds sequencing indexes. Illumina sequencing is performed using 101-bp single end reads, thereby quantifying host mRNA and viral RNA from single cells. (B-C) In total, 127 cells were successfully captured and profiled: 25 Mock cells, 68 WNV cells, and 34 WNV (+Ab) cells. (D) Coverage and alignment of WNV reads are shown with reference to the WNV genome and WNV SC primer (viral primer) location, and y-axes are in log10 scale. The cells representing the median value for WNV and Mock conditions are shown. Violin plot showing expression as counts per million transcripts (CPM) in log2 scale for WNV RNA in all three conditions described in 2A. Wilcoxon rank-sum test with continuity correction was performed to test significance (\*\**p* < 10^−9^). (E) Violin plots showing expression as CPM in log2 scale for housekeeping genes. Wilcoxon rank-sum test with continuity correction was performed to test significance (ns = not significant; \**p* < 10^−3^; \*\**p* < 10^−6^).

Viral RNA was successfully recovered from single cells following WNV infection, and the majority of WNV reads were aligned with the targeted region of the WNV genome (Fig. 2D). To ensure that the addition of WNV SC primer did not adversely affect the recovery of host mRNAs, the concentration of WNV SC primer was carefully titrated and cDNA quality was evaluated on an Agilent 2100 Bioanalyzer (Supplementary Fig. 1). Furthermore, we examined the levels of housekeeping genes (*Gapdh*, *Rpl5*, *Arf1* and *Pgk1*) across cells in all three conditions: Mock, WNV, and WNV (+Ab). Unsurprisingly, expression of housekeeping genes was not significantly different between mock and infected conditions, demonstrating that amplification of viral RNA does not impair recovery of host mRNA (Fig. 2E).

### Heterogeneity in viral RNA abundance and ISG induction at single-cell resolution

At the single-cell level, we observed large differences in viral RNA abundance in the presence and absence of limited *in vitro* spread (Fig. 2D). In the absence of neutralizing antibody, we detected a wide range of intracellular viral RNA levels, with the majority of cells having greater than 2^10^ viral RNA counts per million transcripts (Fig. 2D). Interestingly, only 24% of cells had greater than 2^10^ viral RNA counts per million transcripts when limiting asynchronous secondary infection (Fig. 2D). Furthermore, the heterogeneity of viral RNA abundance in the presence of neutralizing antibody suggests that there is variability in WNV replication during the primary round of infection (Fig. 2D). Notably, the percentage of single cells positive for viral RNA (Fig. 2D) is significantly higher than the percentage predicted by flow cytometry-based viral E protein immunostaining for both infection conditions (Fig. 1B).

When examining transcriptional dynamics across single cells, we noticed some interesting trends. Only a small fraction of WNV-infected cells produced greater than 2^5^ *Ifnb1* counts per million transcripts (Fig. 3A). Intriguingly, we observed a similar expression signature for *Ifna4* and *Ifna2* despite high levels of *Irf7*, a transcription factor that drives IFN-α production (47–49), in the majority of cells (Fig. 3A). Furthermore, we identified three chemokine genes (*Ccl5*, *Ccl4* and *Cxcl11*) that displayed comparable cellular distributions to IFN-I genes. Other pro-inflammatory cytokine genes, *Cxcl10*, *Tnf*, *Il6* and *Il23a*, exhibited cellular heterogeneity but still maintained a portion of cells with no detectable transcript. Genes *Ddx58* and *Dhx58*, which respectively encode the RLRs RIG-I and LGP2 (Laboratory of Genetics and Physiology 2), were highly expressed with most cells containing greater than 2^5^ counts per million transcripts (Fig. 3B). Interestingly, *Tlr3* and *Ifih1*, another important RLR gene that encodes MDA5 (melanoma differentiation-associated gene 5), displayed greater variation in expression across cells, including a fraction with no detectable transcript (Fig. 3B). Components of the ISGF3 complex (*Irf9*, *Stat1* and *Stat2*) are critical for IFN-I signaling and are induced to greater than 2^5^ counts per million transcripts in the majority of cells (Fig. 3B). Next, we sought to examine the expression patterns for a panel of experimentally validated WNV-targeting antiviral effector genes (*Rsad2*, *Tnfsf10*, *Ifi44l*, *Oas1b*, *Oas3*, *Ifitm3*, *Eif2ak2* and *Mov10*) and two well-established ISGs (*Ifit3* and *Mx1*) (26, 42, 50–58). Antiviral effector genes feature both unimodal (*Tnfsf10*, *Ifi44l*, *Ifitm3*, *Eif2ak2*, *Mov10* and *Mx1*) and bimodal (*Rsad2*, *Oas1b*, *Oas3* and *Ifit3*) variation across single cells (Fig. 4). Notably, several genes (*Ddx58*, *Tlr3*, *Stat1*, *Tnfsf10*, *Eif2ak2*, *Ifit3* and *Mx1*) revealed significantly different transcriptional signatures across cells with and without limited *in vitro* spread (Fig. 3B, 4). Strikingly, most cells have no detectable reads for *Tnfsf10* and *Mx1* when allowing for *in vitro* spread; however, in the presence of neutralizing Ab, the inverse is true (Fig. 4).

**Figure 3.**
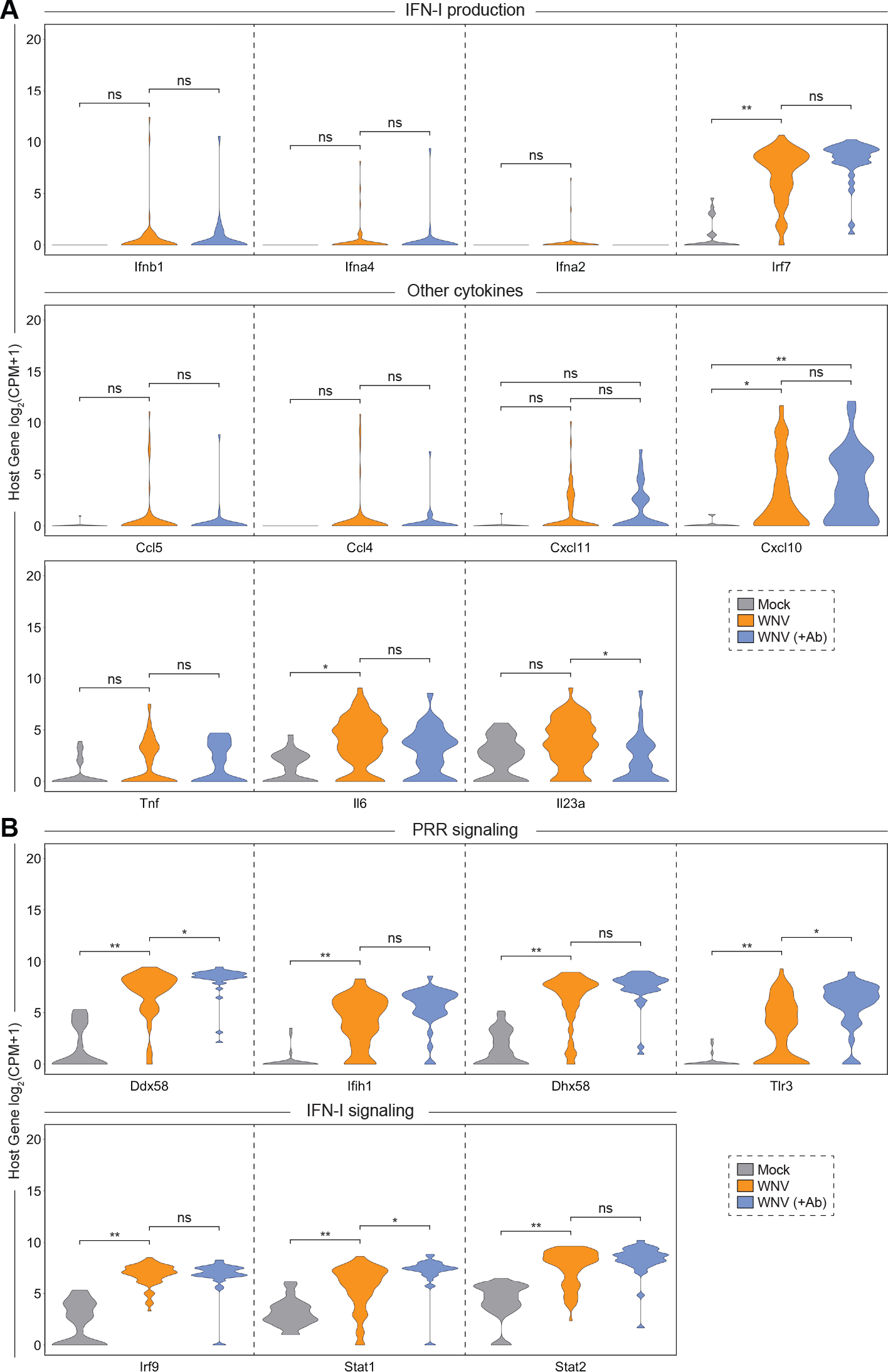
Cellular heterogeneity in IFN-stimulated gene induction following WNV infection. Violin plots showing single-cell distributions for host gene expression as counts per million transcripts (CPM) in log2 scale. Genes are grouped by categories: (A) IFN-I production and other cytokines; and (B) PRR and IFN-I signaling. Conditions are described in Fig. 2A. Wilcoxon rank-sum test with continuity correction was performed to test significance (ns = not significant; \**p* < 10^−3^; \*\**p* < 10^−6^).

**Figure 4.**
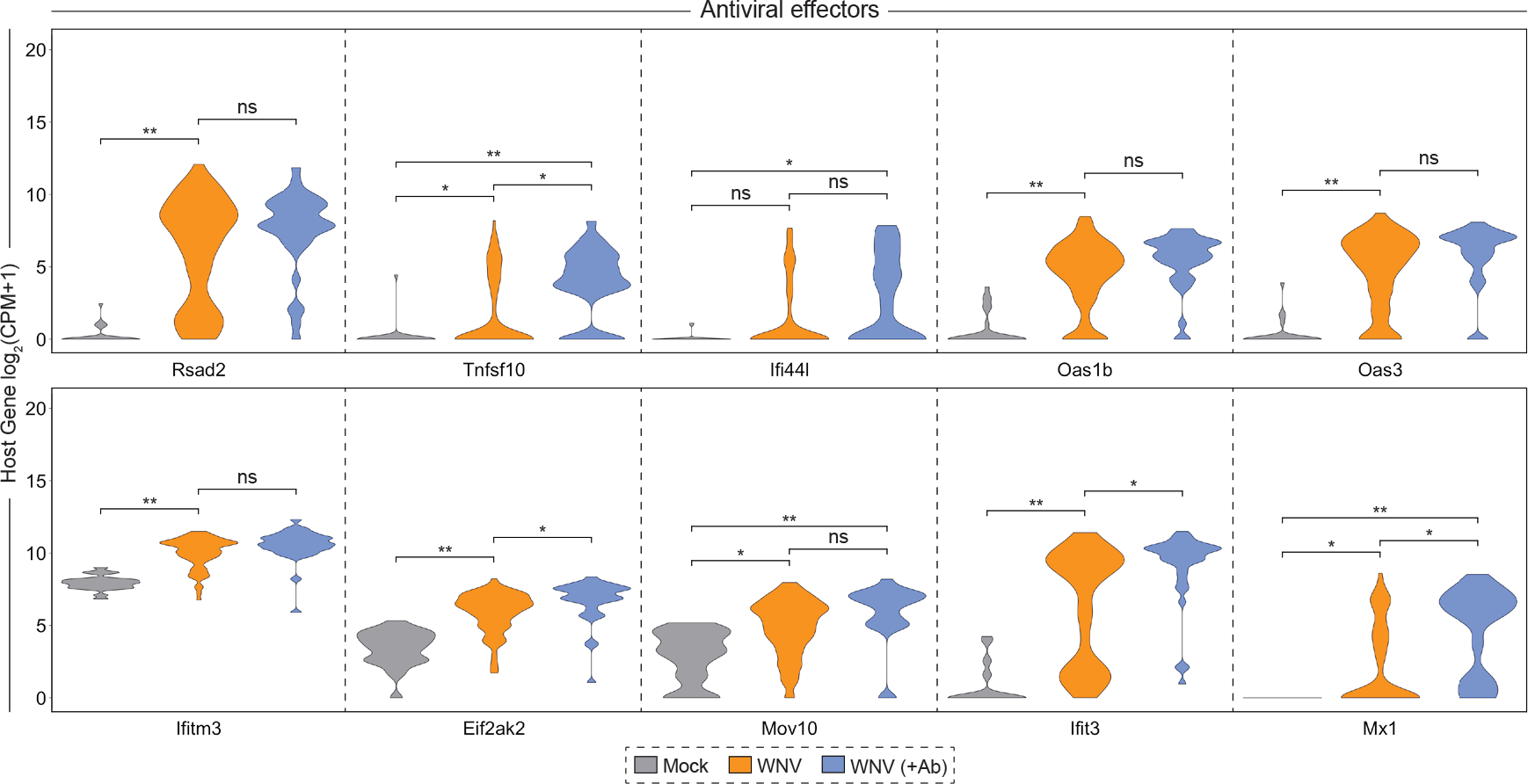
Unimodal and bimodal variation in antiviral effector gene expression at single-cell following WNV infection. Violin plots showing single-cell distributions for antiviral effector gene expression as counts per million transcripts (CPM) in log2 scale. Conditions are described in Fig. 2A. Wilcoxon rank-sum test with continuity correction was performed to test significance (ns=not significant; \**p* < 10^−3^; \*\**p* < 10^−6^).

### Correlation between host gene expression and viral RNA abundance for single cells

Building upon our ability to assess viral RNA abundance in singe cells, we calculated Spearman’s correlation coefficients for host gene expression and viral RNA burden across all WNV cells, which spanned a range of viral RNA levels. To comprehensively identify pathways that might be linked to viral RNA abundance, we performed gene ontology (GO) enrichment analysis using the online bioinformatics tool DAVID (59, 60), wherein we independently evaluated all positively correlated (ρ > 0.35) and negatively correlated (ρ < −0.35) genes of significance (*p* < 0.001). The top pathways extracted from the GO enrichment analysis for negatively correlated genes were the antiviral defense response, cellular response to IFN-β, response to virus, negative regulation of viral replication, innate immune response and antigen processing and presentation via MHC I (class I major histocompatibility complex molecule) (Fig. 5A). For the positively correlated gene set, the top pathways included transcriptional regulation, amino acid transport, ribosomal RNA (rRNA) processing, regulation of protein ubiquitination and ER stress response, providing a broad description of viral RNA-correlated genes (Fig. 5A). Next using curated gene lists from published large-scale ISG screen and single-cell transcriptomic studies (26, 61, 62), we examined the distribution of correlation coefficients for ISGs and cell cycle-associated genes subdivided by phase (G1/S, S, G2/M, M and M/G1). Predictably, the majority of genes do not correlate with viral RNA abundance, and the distribution of coefficients skews heavily towards zero (Fig. 5B). When assessing viral RNA correlations for cell cycle-associated genes, most genes were not significantly positively or negatively correlated, although minor shifts were observed for S, M and M/G1 phase genes (Fig. 5B). Interestingly, 124 out of 294 ISGs were negatively correlated with viral RNA corresponding with a dramatic shift in the coefficient distribution (Fig. 5B). As predicted by the GO enrichment analysis, numerous genes associated with the ER stress response (*Gadd45a*, *Ppp1r15a*, *Selenos*, *Ddit3*, *Atf4* and others) were strongly positively correlated with viral abundance (Fig. 5B). A subset of correlated ISGs and panel of non-correlated cytokine genes are represented in a correlation matrix (Fig. 5C). Negatively correlated ISGs strongly clustered together with high correlation coefficients approaching 1 (Fig. 5C). Conversely, ISGs positively correlated with viral RNA only weakly correlated with other positively correlated ISGs (Fig. 5C). Many cells expressing high levels of IFN-I and pro-inflammatory cytokines also featured elevated viral abundance, but not to the extent of reaching significant positive correlation (Fig. 5C). Scatter plots were generated for a subset of viral RNA-correlated genes and collated in order of increasing correlation coefficients (Fig. 6). Trends associated with negatively correlated ISGs mostly featured a precipitous decline in gene expression as viral RNA levels in single cells exceeded around 2^15^ counts per million transcripts (Fig. 6). Alternatively, positively correlated genes often were characterized by slopes near or less than 1 (Fig. 6). For WNV-validated antiviral effector genes (*Rsad2*, *Tnfsf10*, *Ifi44l*, *Oas1b*, *Oas3*, *Ifitm3*, *Eif2ak2* and *Mov10*), all genes are negatively correlated with viral RNA as expected (Fig. 6). Interestingly, *Tnfsf10*, *Ifi44l* and *Mx1* present unique correlation trends with viral RNA in that the cells with the highest viral abundance have no detectable transcripts for these genes (Fig. 6).

**Figure 5.**
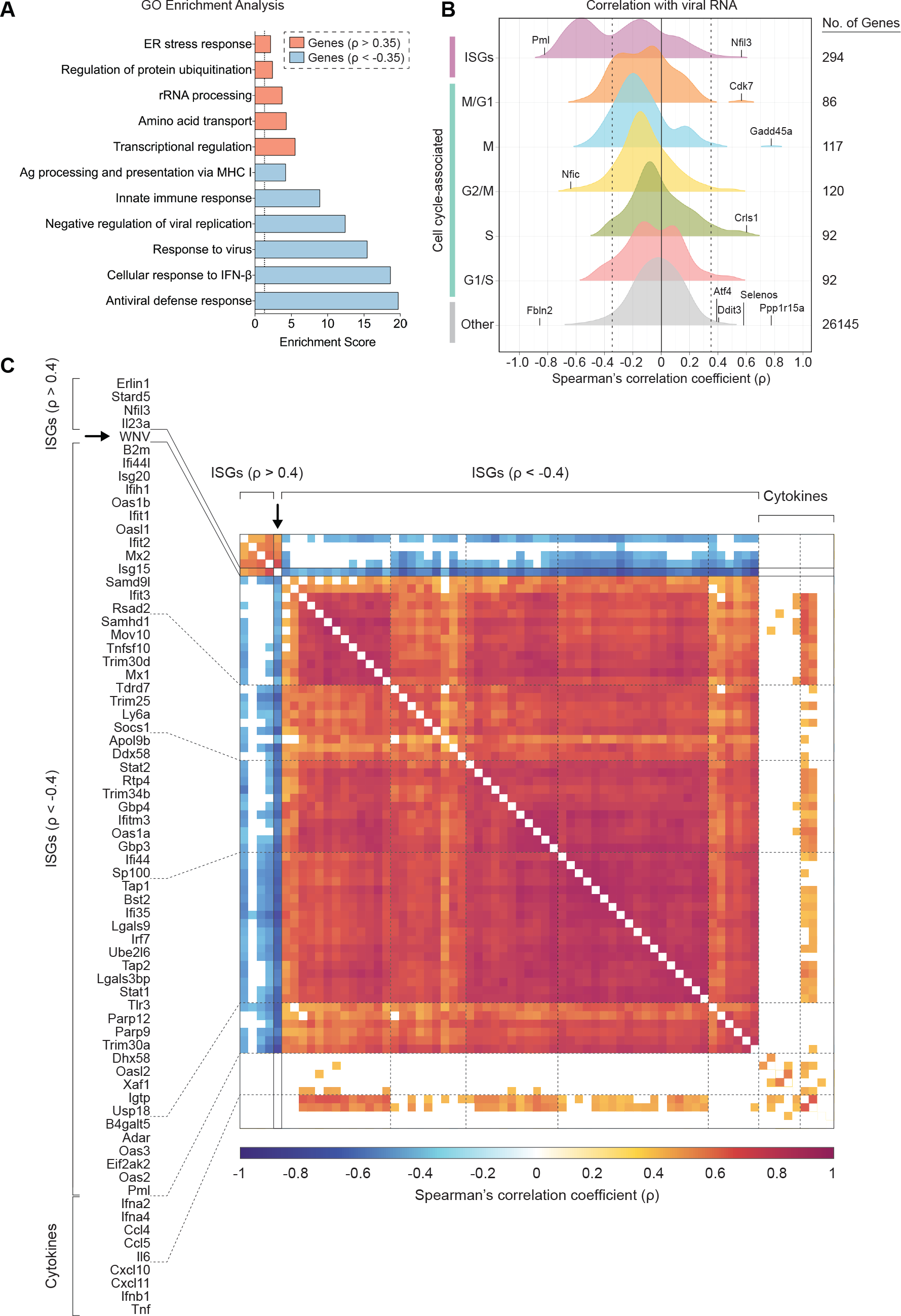
ISGs negatively correlate with WNV RNA abundance. (A) Gene ontology (GO) enrichment analysis for genes significantly (*p* < 0.001) positively correlated (ρ > 0.35) and negatively correlated (ρ < −0.35) with viral RNA. Enrichment scores (ES) calculated for each pathway by the formula: −log_10_(*p* value). Dotted line indicates significance cutoff (*p* = 0.05; ES = 1.3). (B) Density plots of host gene expression correlated viral RNA across all WNV cells. Spearman’s correlation coefficients (ρ) calculated for each host gene by viral RNA. Gene set labels (left) and totals (right) are shown. Cell cycle-associated genes are additionally subdivided by phase. Select genes were marked and labeled. Dotted lines indicate correlation coefficients (ρ) equal to −0.35 and 0.35. (C) Correlation matrix for 57 of 124 negatively correlated ISGs, all positively correlated ISGs, 9 non-correlated cytokine genes and WNV RNA. Correlation coefficients (ρ) calculated for each gene pairing are indicated by the color gradient (bottom). White boxes represent comparisons for which the correlation did not meet the significance cutoff (*p* < 0.001).

**Figure 6.**
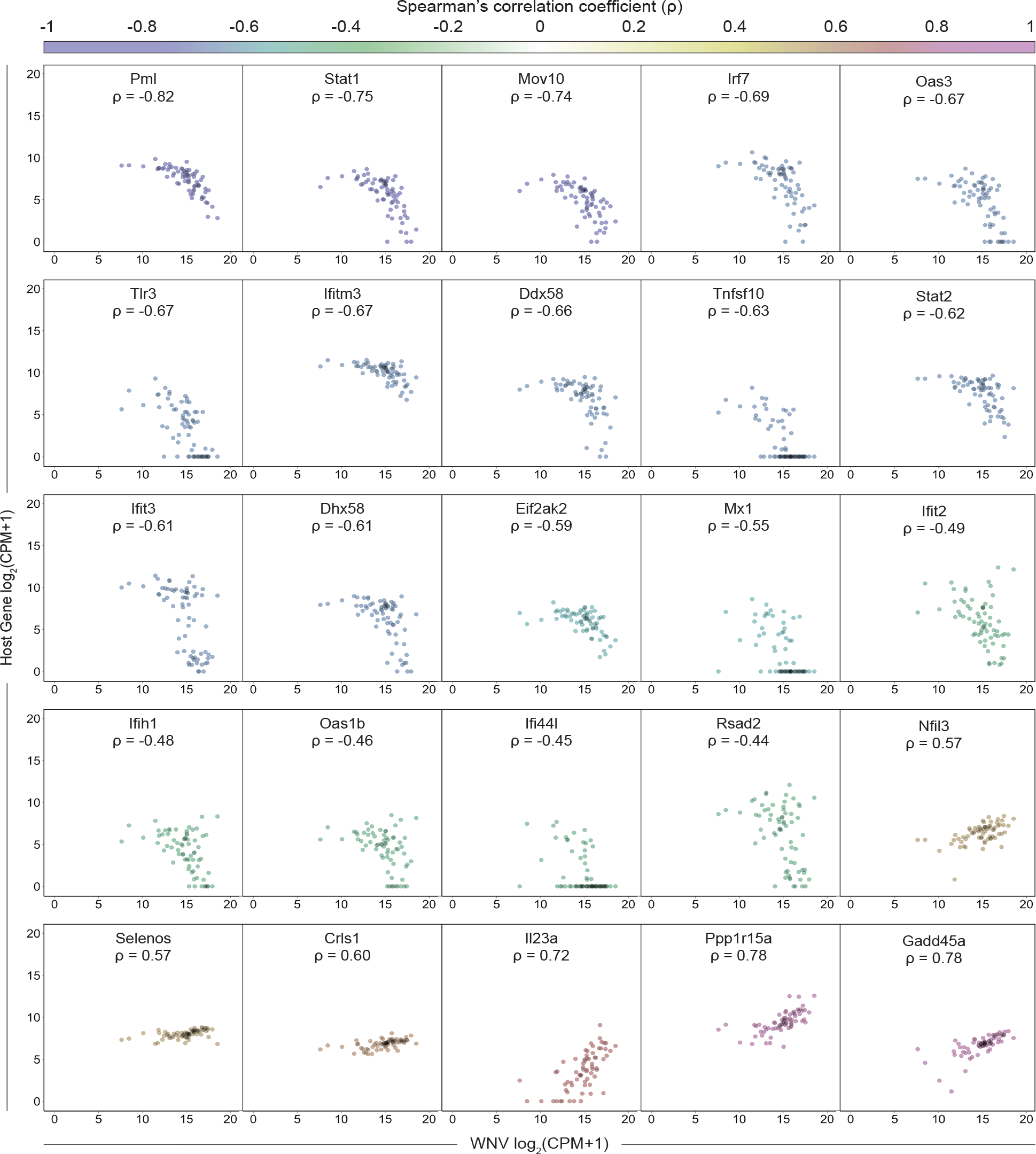
Sharp downward trend for ISGs negatively correlated with viral RNA. Scatter plots showing expression as counts per million transcripts (CPM) in log2 scale of positively and negatively correlated host genes by WNV RNA. Each cell is represented by a single dot with minimal transparency so areas of high density are easily discernable. Correlation coefficients (ρ) are indicated for each gene and correspond to the color gradient (top). Scatter plots have been collated from lowest to highest correlation coefficient. All genes shown here meet the following criteria: |ρ| > 0.4 and *p* < 0.0005.

## DISCUSSION

Standard scRNA-seq protocols with oligo-dT-based priming have been used to examine transcriptional dynamics during viral infection, but the unique genomic structure of flaviviruses, and other non-polyadenylated viruses that are clinically important pathogens, represents a distinct hurdle for such studies (29). We have independently developed and demonstrated the feasibility of WNV-inclusive scRNA-seq as an attractive approach for the quantification of host transcripts and viral RNA within single cells. This protocol, in combination with previously published work by Zanini and colleagues (31), establishes virus-inclusive scRNA-seq as a viable and tractable system for other non-polyadenylated RNA viruses.

WNV-inclusive scRNA-seq revealed extensive transcriptional heterogeneity in viral RNA abundance and the IFN-I response across single cells (Fig. 2D, 3, 4). The majority of WNV reads mapped to the targeted region of the WNV genome (Fig. 2D). However, minimal non-random background was observed in Mock cells with a median value of 45 WNV CPM for Mock cells, as compared to 37246 WNV CPM for WNV cells. This background may result from index hopping (63), and could be accounted for in subsequent iterations by using unique indexes. In support of published findings (17–19, 27), we found that few cells produce IFN-β transcript following viral infection (Fig. 3A). However, we observed a strong induction of numerous ISGs (*Irf7*, *Ddx58*, *Dhx58*, *Irf9*, *Stat1* and *Stat2*) with high unimodal expression signatures (Fig. 3), highlighting the well-established importance of IFN-I-dependent paracrine signaling (16–19, 26, 27). Interestingly, we saw a bifurcation in ISG correlations with viral RNA, wherein 124 out of 294 ISGs were negatively correlated with intracellular viral abundance (Fig. 5B). Furthermore, a considerable fraction of ISGs featured a precipitous downward trend in expression with increasing viral RNA, dissimilar to the gradual upward trend exhibited by positively correlated genes (Fig. 6). Collectively, these findings are reflective of the dynamic balance and interplay between host and viral factors within a single cell. This represents the first single-cell transcriptomics study of flavivirus infection to examine the correlation of ISGs with intracellular viral RNA. To extend this arm of our analysis, we examined WNV-targeting antiviral effector genes that have been previously validated through short hairpin RNA (shRNA) and small interfering RNA (siRNA) knockdown screens, cell-based overexpression assays and *in vivo* knockout models (26, 42, 50–58). These validated antiviral effector genes exhibited both unimodal (*Tnfsf10*, *Ifi44l*, *Ifitm3*, *Eif2ak2* and *Mov10*) and bimodal (*Rsad2*, *Oas1b* and *Oas3*) expression patterns and all negatively correlated with viral RNA (Fig. 4, 6), demonstrating the technical capacity of WNV-inclusive scRNA-seq to probe virus-host interactions and identify novel antiviral candidate genes.

The discovery that bimodal variation in IFN-stimulated genes (ISGs) correlates with viral RNA abundance (Fig. 4, 6) bears notable relevance to previous work examining WNV antagonism of IFN-I signaling. WNV, among other flaviviruses, directly or indirectly antagonizes IFN-I signaling and the JAK-STAT pathway to counter cellular antiviral defenses (64–68). The WNV nonstructural protein NS5 blocks Jak1 and Tyk2 activation by interacting with prolidase to inhibit surface expression of IFNAR1 (10, 64). Additionally, WNV recruits plasma membrane-derived cholesterol to replication sites in the ER, and NS4A and NS4B contribute to membrane rearrangement and associated ER stress, which are all thought to interfere with JAK-STAT signaling (64, 69–71). Bimodal ISG expression patterns correlated with viral abundance (Fig. 4, 6) may result from viral antagonism in primary infected cells allowing for higher replication. This is supported by the almost uniformly high expression observed for ISGs when limiting *in vitro* spread (Fig. 4), a cell population with a predominantly low-level of WNV replication (Fig. 2D). Alternatively, bimodality may arise from preexisting cell-intrinsic differences, such as the level of critical signaling components, specifically at the initial stage of infection.

WNV-inclusive scRNA-seq provides a single-cell transcriptomics protocol to probe cellular heterogeneity in the host response and quantify viral RNA. The outlined approach can potentially serve as a valuable tool for *in vivo* studies to examine cell-intrinsic responses to viral infection, extending the resolution to infected single cells. Such studies can also leverage the added ability with this approach to screen for infected cells by qPCR and cherry pick cDNA for sequencing to mitigate cost. Our study demonstrates the feasibility and utility of WNV-inclusive scRNA-seq as a high-throughput technique for single-cell transcriptomics and viral RNA detection, which can be used to provide insights into the cellular features of protective immunity.

## MATERIALS AND METHODS

### Cells and viruses

Murine fibroblast L929 cells were obtained from ATCC and grown at 37°C with 5% CO_2_ in DMEM (Corning) supplemented with 10% heat-inactivated FBS, 2mM L-glutamine (Corning), 25 mM HEPES buffer (Corning), 1mM sodium pyruvate (Corning), MEM nonessential amino acids (Corning) and antibiotics/antimycotics (Corning). WNV isolate Texas 2002-HC (WNV-TX) has been previously described (3, 72, 73), and its titer was determined by standard plaque assay on BHK-21 cells. Working stocks were generated by passaging WNV-TX twice on Vero cells (ATCC CCL81) and used for *in vitro* experiments. WNV was incubated directly under ultraviolet (UV) light for 1 hr to generate UV-inactivated WNV, which was confirmed by plaque assay prior to use.

### Infection and antibody treatment

L929 cells were plated to 70-80% confluent and infected with WNV-TX at different MOIs (0.1, 1 or 10). Following a 1 hr virus adsorption period at 37°C, cells were washed once with complete DMEM (cDMEM) and subsequently incubated for 6-48 hr with cDMEM or cDMEM supplemented with WNV E16 neutralizing antibody (5 μg/mL), a gift from Michael Diamond (Washington University, St. Louis, Missouri) (44). Cells were trypsinized for flow staining or lysed for RNA at 6, 12, 24 or 48 hr post-infection. Antibody titration in supplemental media was performed at multiple MOIs (0.1, 1 or 10) for 48 hr post-infection to determine the optimal concentration to considerably reduce *in vitro* spread prior to use.

### Flow cytometry

Conditions were run in biological triplicate samples. Cells were treated with 0.125% trypsin in PBS for 5 min at 37°C. All centrifugation steps were performed at 1250 rotations per minute for 5 min at 4°C. Cells were pelleted, resuspended in FACS buffer (1x PBS, 1% FBS, 1 mM EDTA), and blocked for 10 min on ice with anti-mouse Fc Shield (TONBO Biosciences) at 0.5 μL per sample in FACS buffer. Subsequently, samples were stained for 20 min on ice with Ghost 780 viability dye (TONBO Biosciences) at 0.1 μL per sample in PBS. Samples were washed and resuspended in FACS buffer. For WNV E protein staining, samples were fixed following viability staining with 1x Transcription Factor Fix/Perm (diluted in Transcription Factor Fix/Perm Diluent; TONBO Biosciences) for 20 min on ice and washed twice with 1x Flow Cytometry Perm Buffer (diluted in ddH_2_O; TONBO Biosciences). WNV E16 antibody was conjugated to Allophycocyanin (APC) using the Lightning-Link APC Antibody Labeling Kit (Novus Biologicals). Samples were stained with APC-conjugated WNV E16 antibody at 0.25 μg per sample in Flow Cytometry Perm Buffer for 30 min on ice. Samples were washed, resuspended in FACS buffer, and run on a BD LSR II flow cytometer.

### Single-cell sorting

Cells were stained with Ghost 780 viability dye (TONBO Biosciences) as stated above and filtered through a 35 μm strainer into a 5 mL FACS tube. Single viable cells were sorted into skirted 96-well PCR plates containing 10 μL RLT buffer (Qiagen) with 2-betamercaptoethanol (1:100) per well using a BSL-3 level BD Aria II flow cytometer.

### Quantitative RT-PCR (qPCR)

Time-matched mock and WNV-infected L929 cells (1 × 10^5^ cells per condition; in triplicate) were lysed in RNA Lysis Buffer. Total RNA was isolated from cells using the Quick-RNA MiniPrep Kit (Zymo Research). Purified RNA was reverse transcribed using random primers with the High-Capacity cDNA Reverse Transcription Kit (Applied Biosystems). WNV RNA levels were quantified by qPCR using PrimeTime Gene Expression Master Mix (Integrated DNA Technologies), WNV-specific primers and probe set, and TaqMan gene expression assay (ThermoFisher) for the host gene *Gapdh* (Mm99999915_g1). WNV-specific primer and probe sequences (Forward primer: 5’ – TCAGCGATCTCTCCACCAAAG – 3’; Reverse primer: 5’ – GGGTCAGCACGTTTGTCATTG – 3’; and Probe: 5’ – 6FAM-TGCCCGACC-ATGGGAGAAGCTC-MGB – 3’) were adapted from Lanciotti and colleagues (73) and correspond to WNV isolate Texas 2002-HC (GenBank accession number: DQ176637.1). C_T_ values were normalized to the reference gene *Gapdh* and represented as fold change over time-matched mock values using the formula 2^−ΔΔCT^. All primers and probes were purchased from Integrated DNA Technologies (IDT). qPCR was performed in 384-well plates and run on an Applied Biosystems 7900 HT Real-Time PCR System.

### Bulk mRNA sequencing (RNA-seq)

L929 cells were infected with WNV (MOI of 1) and incubated in the absence or presence (+Ab) of WNV E16 neutralizing Ab. In biological triplicate (n = 3), 50,000 viable cells were sorted into 100 μL RLT buffer (Qiagen) with 2-betamercaptoethanol (1:100) at 24 hr post-infection for each condition: time-matched mock, WNV and WNV (+Ab). mRNA sequencing libraries were prepared at Yerkes Genomics Core (http://www.yerkes.emory.edu/nhp_genomics_core/), and the quality of the libraries was verified using DNA-1000 Kits (Agilent Bioanalyzer) and quantified using the Qubit 2.0 Fluorometer (LifeTechnologies). Libraries were clustered and sequenced on an Illumina HiSeq (100 bp single-end reads). Sequencing reads were mapped to the GENCODE mouse reference genome (GRCm38.p5 Release M16). Reads were normalized and differential expression analysis performed using DESeq2 (74). Normalized reads were expressed as fold change over time-matched mock values.

### Single-cell RNA sequencing (scRNA-seq)

SMART-Seq v4 Ultra Low Input RNA Kit (Takara) was used for cDNA preparation. The protocol was modified to include a WNV-specific viral primer during the RT step. WNV SC primer (5’ – <u>AAGCAGTGGTATCAACGCAGAGTAC</u>GGGTCAGCACGTTTGTCATTG – 3’) targets the positive-sense envelope protein (E) gene (73) and contains the 5’ 25-nucleotide ISPCR universal anchor sequence (underlined) from the Smart-seq2 protocol published by Picelli and colleagues (45) for downstream amplification alongside 3’ SMART-Seq CDS Primer II A-primed transcripts. Similar to 3’ SMART-Seq CDS Primer II A, 1 μL of WNV SC primer (12 μM) was added to the RT reaction for all samples. Other WNV-specific primer sequences and concentrations were evaluated. The scRNA-seq protocol was optimized to ensure high sensitivity for WNV RNA detection and to mitigate the formation of primer dimers or template-switching oligo (TSO) concatemers observed at high concentrations or with other primer sequences. During template switching, the RT product is extended with a sequence complementary to the TSO due to the addition of 2-5 untemplated nucleotides and the capacity of the RT enzyme to switch templates just as described in Smart-seq2 (45). PCR is performed using PCR Primer II A (the ISPCR universal anchor sequence) for concurrent amplification of both host and viral cDNA. Following PCR amplification, cDNA quantification is performed for each sample, and cDNA quality assessment is accomplished using an Agilent 2100 Bioanalyzer. For library preparation, amplified cDNA is fragmented and appended with dual-indexed barcodes using Illumina Nextera XT DNA Library Prep kits. Sequencing was performed using 101-bp single end reads at Yerkes Genomics Core (http://www.yerkes.emory.edu/nhp_genomics_core/) as previously described (75) on an Illumina HiSeq 3000 at a depth of ∼1,000,000 reads per cell. In total, 127 cells were successfully captured and profiled for single-cell transcriptomic analysis: 25 Mock cells, 68 WNV cells, and 34 WNV (+Ab) cells.

### Bioinformatics pipeline

Libraries were sequenced on an Illumina HiSeq 3000 generating 101-bp single end reads. FastQC (76) was used to check the quality of fastq files. The primary assembly of GENCODE mouse reference genome (GRCm38.p5 Release M16) (77) and the complete genome of WNV isolate Texas 2002-HC (GenBank accession number: DQ176637.1) from ViPR (78) were used for mapping reads. The genome index was built by combining both the genomes, and alignments were carried out for the combined genomes. STAR v2.5.2b (79) was used with default parameters to map reads and obtain reads per gene counts (–quantMode Gene Counts). The counts obtained with STAR were used for downstream analysis in R. The counts were used to create a SingleCellExperiment v1.0.0 (80) object. The scater v1.6.3 (81) library was used for quality control of cells. Genes that were not expressed in any cell were filtered out. The isOutlier function from scran was used to remove cells that had a library size and number of detected genes greater than 3 median absolute deviations lower than the median values or those with percentage of mitochondrial genes that were 3 median absolute deviations higher than the median value (82). The cell cycle phase was predicted using the cyclone function in scran package v1.6.3 (80, 82). The normalized expression values were obtained using the calculateCPM function in the scater library.

### Statistical analysis and software

Prism 6 (GraphPad), ggplot2 R package, ggridges R package, corrplot R package and Hmisc R package were used for statistical analyses and graphical presentation of data. Spearman’s rank correlation coefficients (ρ) and associated *p* values were computed for each gene pairing using the rcorr function in Hmisc R package. Two-way ANOVA with Tukey’s multiple comparison correction was used to evaluate significant differences between conditions for percentage of WNV E-positive cells and relative viral RNA. Wilcoxon rank-sum test with continuity correction was performed to assess significant differences between single-cell distributions for host mRNA and viral RNA counts per million transcripts (CPM).

## Acknowledgments

We thank the Emory Vaccine Center Flow Core, specifically Kiran Gill and Barbara Cervasi, for assistance with cell sorting and the Yerkes Genomics Core for library preparation and sequencing.

